# Elucidating an implicit-statistical learning brain network: Coordinate-based meta-analyses and functional connectivity profiles of artificial grammar learning in healthy adults

**DOI:** 10.1101/2024.01.10.575023

**Authors:** Amy E. Ramage, Kaila Cote, Jill C. Thorson, Katelyn Lerner, Michael C. Reidel, Angela R. Laird

## Abstract

Language rehabilitation centers on modifying its use through experience-based neuroplasticity. Implicit statistical learning of language is essential to its acquisition and likely its rehabilitation following brain injury, but its corresponding brain networks remain elusive. Coordinate-based meta-analyses were conducted to identify common and distinct brain activity across 25 studies coded for meta-data and experimental contrasts (grammatical or non-grammatical). The resultant brain regions served as seeds for profiling functional connectivity in large task-independent and task-dependent data sets. Hierarchical clustering of these profiles grouped brain regions into three subnetworks associated with grammatical/non-grammatical processes. Functional decoding clarified the mental operations associated with those subnetworks. Results support a left-dominant language sub-network and two cognitive control networks as scaffolds for grammar rule identification, maintenance, and application in healthy adults. These data suggest that cognitive control is necessary to track regularities across stimuli and imperative for rule identification and application of grammar. Future empirical investigation of these brain networks in language learning in individuals with brain injury will clarify their prognostic role in language recovery.

## 1. Introduction

Rehabilitation for cognitive-communication impairments following left hemisphere stroke requires *re-*learning, as behavioral interventions train restorative or compensatory skills to improve performance. However, little is known about how the damaged brain *re*-learns. There are favorable brain-based biomarkers of recovery from stroke (Basso, 1992; Busby et al., 2023; Fedorenko et al., 2013; Gilmore et al., 2018; Hartwigsen et al., 2013; Hope et al., 2017; Ramage et al., 2020; Saur et al., 2006), but not as much is known about how rehabilitation efforts lead to or enhance brain changes to result in favorable recovery of communication skills. Thus, we explore a return to the idea of Baddeley (1993), and posit that a *model of learning* is needed to inform how we approach rehabilitation. Utilizing such a model will help to optimize interventions by leveraging changes in brain activity or connectivity to attain positive outcomes.

In a first step to investigating models of learning for rehabilitation of language, we identify language learning paradigms and their corresponding brain networks in neurotypical, healthy adults. We focus on learning that is known to be involved in language acquisition, i.e., implicit or statistical learning (c.f., Perruchet & Pacton, 2006) because this process is also utilized in adult language learning. Most often, implicit, or statistical language learning is investigated through artificial grammar learning in experimental paradigms in which participants are exposed to novel language-like stimuli and must delineate salient cues, or rules, in incoming stimuli without explicit instruction to do so. The learner may receive reinforcement when that salient stimulus is paired to a relevant response, but do not otherwise receive explicit feedback. Individuals with aphasia and traumatic brain injury also evidence this type of learning in implicit learning paradigms (Cope et al., 2017; Peñaloza et al., 2015, 2017; Schuchard et al., 2017; Schuchard & Thompson, 2017; Vallila-Rohter & Kiran, 2013a, 2013b), suggesting that it relies on brain systems that differ from those underlying the language impairment. However, there is not consensus on the brain regions or networks associated with implicit learning of language-like stimuli. Modeling implicit learning in the brain in healthy adults can provide *a priori* hypotheses for investigating *re*learning in individuals with acquired brain injury. With such a brain-based model, investigation may commence to determine whether and how an implicit learning brain network is impaired or may be optimized in response to intervention.

### 1.1 Implicit-Statistical Learning (ISL)

Implicit-statistical learning (ISL) is the ability to learn the lawfulness of sequenced sounds, characters, or linguistic units to efficiently respond to those stimuli without explicit knowledge of the rules or conscious strategy use. This definition is an evolution of the implicit and statistical learning literature as follows. *Implicit learning* is the recognition and integration of regularities encountered in the world that occurs without an intention. It is the passive acquisition of knowledge without awareness of what has been learned. ISL is typically measured experimentally utilizing tasks that provide no explicit instruction, and yet an increase in accuracy implies that a participant has implicitly learned the rules of the task (Reber, 1967).

*Statistical learning* is more narrowly defined as the ability to detect transitional or distributed dependencies among elements in close temporal or spatial proximity of each other in a perceived stream of input (Christiansen, 2019; Perruchet & Pacton, 2006). Statistical learning has been investigated in language learning because individuals are known to utilize statistical regularities (e.g., predictive dependencies in word segmentation (Saffran, 2001)) when learning to extract phrase structures, or syntax, that are organized hierarchically. Phrases are marked by dependencies – i.e., one word class depends on another specific word class to follow it. For example, determiners such as “a” or “the” must be followed by a noun. However, the phrases and their constituent probabilities are hierarchically organized in language such that they do not happen bidirectionally (e.g., a noun does not require a determiner) and thus are not based simply on co-occurrence. An example of a transitional probability may be: given B, what is the likelihood of A? Or, given a determiner, what is the likelihood that a noun will follow? Learning of transitional probabilities is evident when learners can detect phrasal units using predictive dependencies above chance, even when all surface level cues (e.g., intonation or lexical stress) are removed (Saffran, 2001; Saffran et al., 1996). The statistical dependencies between the word classes (e.g., nouns and verbs, or nouns and determiners) are as complex as surface level cues and provide a framework for the complex syntactic rules that can be implicitly learned from natural language. Regardless of the considerable spectrum of complexity that may be explained by recognizing statistical dependencies, statistical learning may be considered a subtype of implicit learning as both are grounded in the basic processes of attending to salient cues, learning, and memory, with an uncontroversial overlap through chunking (Christiansen, 2019). That is, implicit learning typically involves rule application is chunked once acquired, and is more efficient when employed for computation in the moment (Batterink et al., 2015).

Therefore, this study considers statistical learning and implicit learning synonymously as implicit-statistical learning (ISL).

### 1.2 Artificial or Non-Native Grammar Learning

While ISL has been studied using several experimental paradigms, artificial or non-native grammar tasks most closely mimic learning of natural language as they require ISL of hierarchical rule learning. A person is exposed to a *language* that requires segmentation of sound strings into words and hierarchical organization of *words* based on implicitly learned rules. Thus, learning requires the recognition of regularity in the input and subsequent extraction of rules from the stimuli. Adults have been shown to reliably extract regularities in unfamiliar languages or artificial grammars, allowing them to make judgements about the grammaticality of an utterance and perform above chance (Bahlmann et al., 2008; Batterink et al., 2015; Lieberman et al., 2004; Petersson et al., 2004, 2012; Plante et al., 2015; Reber, 1967). Artificial or non-najve grammar tasks are composed of learning and testing phases. In the learning phase, participants are exposed to multiple repetitions of grammatical stimuli demonstrating permissible sentence composition in the grammar. Participants may or may not receive instructions for the task, but the rules are never explicitly taught. Participants then enter the testing phase during which they make grammaticality judgements about presented grammatical sentences, random word-ordered sentences, or sentences that partially follow a rule. Participants can efficiently and accurately judge the grammaticality of sentences created from an artificial grammar while being unaware of the underlying rules or of strategies they may have used.

### 1.3 Neural Models of Language Learning through ISL

Differing lines of research exist that contribute to understanding how the brain accomplishes ISL. Some models center on domain-general or modality-specific processes necessary for different phases of learning. For example, Frost, Siegelman, Christiansen and colleagues (Christiansen, 2019; Christiansen et al., 2010; Christiansen & Chater, 2016; Frost et al., 2015; Siegelman et al., 2017, 2018; Siegelman & Frost, 2015) highlight the similarities across implicit learning tasks, i.e., that learning requires extraction of salient features in stimuli that may be patterned/rule-governed and integration of those patterns/rules to apply to other stimuli. In a review of neuroimaging findings in ISL tasks, Frost and colleagues propose unimodal brain regions are dedicated to processing of modality-specific information (e.g., occipital cortex for visual stimuli, temporal cortex for auditory stimuli) and recognize statistical regularities to a greater extent than random information in the stimuli (Frost et al., 2015). However, there also are brain regions involved in ISL regardless of stimulus modality, that may be considered domain-general. These regions include the hippocampus, basal ganglia, thalamus, and prefrontal cortex that are thought to operationalize or modulate modality-specific information.

Ullman and colleagues have taken a different tack by focusing on the type of memory, or memory system, engaged in learning language. The Declarative-Procedural Model (DPM, (Ullman, 2004)) focuses specifically on language learning and the type of linguistic information being learned (rule-governed or lexical). The DPM distinguishes a mental lexicon that is learned through explicit means and stored as part of declarative memory, and a mental grammar learned through implicit means and stored in procedural memory. The *mental lexicon* is a storage of all memorized word-specific knowledge. This includes knowledge of word meanings and sounds, but also any language unit that cannot be derived from another, such as inflectional bound morphemes or idiomatic phrases. The mental lexicon is responsible for the encoding, storage, and use of semantic and episodic memory for facts and events. Information in the mental lexicon is learned explicitly, can be explicitly recalled, and is proposed to be subserved by the declarative memory system. This memory system involves the medial temporal lobes (hippocampus, dentate gyrus and subicular complex, parahippocampus, entorhinal and perirhinal cortex) for formation and consolidation of new memories (c.f., Winocur et al., 2010), and distributed association cortex for long-term semantic and episodic memory storage (Batterink & Paller, 2019; Poldrack et al., 2001).

Conversely, the *mental grammar* is a computational system that extracts regularities from language, and analyzes it based on knowledge of rules and constraints. It is part of the procedural memory system, which is responsible for learning new skills and controlling established skills, habits, and procedures. Procedural (implicit) memory utilizes brain regions similar or related to those for declarative memory, but also involves brain structures historically associated with motor skill learning (a frontal/basal ganglia network) or with conflict resolution/prediction of behavior (superior temporal lobe, parietal lobe, and cerebellum).

Functionally, the basal ganglia are thought to be involved in implicit learning generally, but are specifically engaged for probabilistic rule learning, sequence learning, context-dependent rule selection, working memory maintenance, and attention shifting (Poldrack et al., 2001; Ullman, 2004). The striatal regions (caudate nucleus and putamen) are particularly involved in explicit and implicit language learning (Copland et al., 2021), but their connections with cortical regions appear to be gated based on the phase of learning. For example, Plante et al. (2015) found that activation in the right caudate nucleus is present immediately preceding behaviorally evidenced learning during a non-native grammar learning task, but this caudate activity is reduced once the task becomes more familiar. Similar findings have been reported in other artificial grammar learning studies (Bahlmann et al., 2008; Forkstam et al., 2006; Lieberman et al., 2004). Within the procedural memory system, the learning and knowledge of procedures is unconscious or implicit. More generally, this system is responsible for the learning and processing of sequences and rule relations in context and is used to process complex linguistic rules such as derivational morphology and syntax. Thus, the DPM distinguishes between two types of language knowledge as explicit versus implicit. The declarative mental lexicon subserves rapid learning following one stimulus presentation, which binds and associates with arbitrarily related information. The procedural mental grammar evolves gradually given several presentations of stimuli, tends to be stable with limited flexibility, and exhibits largely unconscious or automatic application of the learned rules.

Specific to artificial grammar or non-native language learning, one region thought to be crucial is the left inferior frontal gyrus (Tagarelli et al., 2019; Yang & Li, 2012). This region appears responsible for hierarchical sequence learning of linguistic *and* non-linguistic stimuli (Forkstam et al., 2006; Karuza et al., 2013; Petersson et al., 2012; Ullman, 2004). In fact, Tagarelli and colleagues, through a series of coordinate-based meta-analyses contrasting grammar learning tasks, found the bilateral IFG involved in language learning for both lexical learning (i.e., word learning in the mental lexicon) and grammar learning. While the left IFG is typically recognized for its involvement in speech and language production (i.e., Broca’s area), it may also serve a role more generally in learning and processing of rule-based sequences. It is also recognized for its role in maintaining information in working memory (Rogalsky et al., 2008). Thus, it is likely that the IFG’s role in working memory supports sequence learning by maintaining information to allow for its consideration or manipulation, which may be essential for the extraction and computation while learning of underlying rules (Arciuli, 2017; Thiessen et al., 2012).

This latter proposal, that the left IFG’s role in grammar learning may be for the general processing of rule-based sequences rather than specifically for language, is in line with Frost and colleagues’ assertion that domain-general regions must be involved in ISL regardless of the modality of the input. Along with the IFG, Tagarelli and colleagues (2019) found activity in a diffuse set of regions, including the premotor and supplementary motor cortex, anterior insula, as well as left-sided superior parietal lobule and angular gyrus during artificial grammar learning tasks whether the intent was to learn word meanings or to learn grammar. These brain regions are also considered part of the dorsal attention network (Corbetta & Shulman, 2002) and salience network (Seeley et al., 2007), along with the cerebellum. For example, the superior parietal lobe, inferior parietal lobe, and supramarginal gyrus, along with the anterior cingulate cortex are engaged in non-native grammar learning tasks in individuals with higher rates of correct rejection of ungrammatical utterances indicating more efficient learning (Plante et al., 2015). As well, the procedural memory system utilizes the cerebellar hemispheres (the dentate nucleus and the vermis) for error-based learning and error detection, two important aspects of grammaticality judgements in ISL tasks (Ullman, 2004).

Thus, the brain activity engaged during artificial and non-native grammar learning tasks and other ISL tasks appears to converge on domain-general and modality-specific regions. Some evidence exists that certain brain systems may be engaged to a greater extent for specific features of the language (i.e., lexical learning = medial temporal cortex; grammar learning = basal ganglia (Tagarelli et al., 2019)). However, it may be that the inter-relationships between the regions is critical to understanding their roles or the processes necessary for ISL of language. When both the declarative (mental lexicon) and procedural systems (mental grammar) are intact, they complement each other. However, when one system is impaired, it may be compensated for by the enhanced function of the other (Ullman, 2004). This is relevant to individuals with acquired brain injury as, for example, damage to Broca’s area (i.e., the left posterior inferior frontal gyrus) results in agrammatic verbal expression reflecting damage to the regions underlying the mental grammar. In contrast, fluent aphasias resulting from lesions in posterior regions of the left hemisphere reflects damage to the medial temporal regions underlying the mental lexicon (Ullman et al., 2005).

The DP model is limited in its explanation of hierarchical rule learning, which relies on recognition of transitional probabilities. Prior to the framework of transitional probabilities, much of the study of statistical learning focused on discovering word segmentation through surface properties of the stimulus, such as the frequency of phonemes or presence of unique units (e.g., distributional statistical learning and prosody). Similarly, artificial grammar learning tasks focus on tracking surface cues such as the frequency of consecutive units (words, syllables, or sounds), frequency of beginning or ending units, lawfulness of the first unit, presence of unique units, location of familiar units, repetition of units, or similarity to previously learned stimuli (Saffran, 2001). However, phrases are not linearly organized in natural language, rather they are more complex (hierarchical), and require more than surface-level cues to extract regularities. Hierarchical organization of phrase structure refers to the organization and grouping of word classes into units (Thompson & Newport, 2007). For example, an English sentence is made up of a noun phrase and a verb phrase, which can also be broken down into smaller groups and categories of words (e.g., determiners, nouns, verbs).

Ullman and colleagues have revised their model to note that some aspects of grammar learning rely on declarative memory, particularly that which is intertwined with lexical learning. Tagarelli et al., 2019 refer to this as *declarative grammar learning* and clarify that this is learning that involves explicit training of grammatical rules, evidence of explicit knowledge of the rules by participants, or when the similarity amongst elements of a grammar is part of the experimental design. In these cases, grammar processing is reliant on declarative memory. They contrast this with *non-declarative grammar learning* from paradigms that more closely align with implicit learning, i.e., when the rules are not trained, and participants are not overtly aware of them.

They found ventral occipito-temporal activation specific to lexical learning, with activity in left inferior temporal and fusiform gyri, but not in the hippocampus, consistent with the “what” or ventral stream (Hickok & Poeppel, 2007). For declarative and non-declarative grammar learning tasks, they found basal ganglia activity (specifically the left caudate head/body and anterior putamen), which they attribute to the role of these structures in procedural memory and automatization of grammar skill. Thus, the Tagarelli meta-analyses largely support the DP model, that explicit and implicit learning is evident in artificial grammar tasks, but also point to involvement of domain-general regions.

The aim of this study is to conduct a coordinate-based meta-analysis of artificial and non-native grammar learning in healthy adults. Specifically, this study will further extend those meta-analytical findings to model functional connectivity profiles and to characterize them with behavioral decoding. There is not currently a meta-analysis that clarifies which brain regions are involved in ISL of grammars, although two meta-analyses have examined language learning (Tagarelli et al., 2019) and non-linguistic sequence learning in adults (Janacsek et al., 2020). While Tagarelli et al. (2019) examined artificial grammar learning in adults, their goal was to look at language learning in general and evaluate the DP model for distinctive regional activity associated with lexical and grammatical learning. As a result, their meta-analysis excluded evaluation of transitional probabilities given that they rely on overlapping lexical and grammatical elements. Nonetheless, their data supported the DP model indicating declarative memory structure involvement in explicit language training and procedural memory structure involvement in implicit training. A focus on the artificial and non-artificial grammars included in their analyses highlighted considerable overlap in the left inferior frontal gyrus for both.

Therefore, the present study will extend the findings of Tagarelli et al. (2019) by identifying the brain regions involved specifically in ISL during artificial and non-native grammar learning tasks. Further, the regions commonly active across studies in the meta-analysis will be used to establish functional connectivity profiles through leveraging of task-independent and task-dependent MRI datasets. These connectivity profiles will be grouped into subnetworks and associated with mental operations. This process will clarify not only the regions uniquely involved in ISL of grammar, but also how those regions integrate with others that are aligned with memory or cognitive systems. We hypothesize that the regions identified as being essential for ISL will be like those reported by Tagarelli et al. (2019). However, we further hypothesize that the evaluation of the functional connectivity profiles will clarify the domain-general networks, outside of the traditional language network, that are engaged.

## 2. Material and Methods

An overview of the methods utilized in the study is presented in Figure 1.

**Figure 1.**
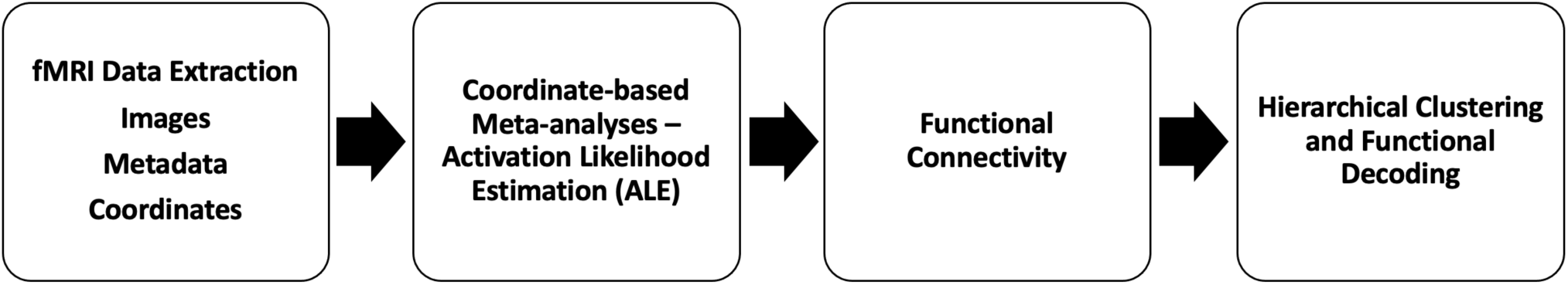
A flowchart demonstrating the steps of the methodological approach. The first step was to evaluate the extant literature involving healthy adults performing ISL tasks in the scanner. As shown in Figure 2, experimental papers were scrutinized for inclusion and exclusion criteria, then the image coordinates and metadata were extracted for entry into the meta-analysis. In the second step, the data were categorized into activation likelihood estimation (ALE) groups to quantify the common brain activity across studies in ISL tasks for either the overall task activity (pooled) or activity elicited when evaluating grammatical or ungrammatical stimuli, as depicted in Figure 3. The ALEs identified regions of interest that were then further evaluated in step 3 to quantify functional connectivity amongst the ROIs by using them as seeds in task-independent (resting-state functional connectivity) and task-dependent functional (meta-analytic connectivity modeling) connectivity analyses as shown in Figure 4. Finally, in step 4 cross-correlations of the *unthresholded* MACMs and rsFC map for each ROI generated correlation matrices that were then averaged. Hierarchical clustering was performed on each matrix. Hierarchical clustering of the connectivity profiles grouped brain regions into subnetworks associated with processes common across spatial and functional domains, as shown in Figure 5.

### 2.1. Literature search, filtering, and paper selection

A comprehensive literature search for peer-reviewed articles commenced from March 27, 2020, to February 19, 2021. PubMed, PsychInfo, Google Scholar, and references of related articles were systematically searched. Additionally, article data from Tagarelli et al. (2019) provided by Ullman (personal communication, 2020) were searched to determine inclusion eligibility for the current study. Inclusion criteria for the studies were: study of healthy adults 18+ years, use of implicit language learning tasks, reporting of coordinates for whole brain analyses, and reporting of experimental contrasts indicating rule learning. Search terms used to find relevant articles included: “implicit learning,” “implicit learning AND automaticity,” “implicit learning AND plasticity,” “implicit learning AND fMRI NOT disorder,” “implicit learning AND statistical learning AND MRI,” “artificial grammar,” “artificial grammar AND fMRI,” “statistical learning AND fMRI,” “artificial grammar learning AND fMRI,” “implicit learning grammar AND fMRI,” and “distributional learning AND fMRI.” Searches yielded over 800 papers. Titles and abstracts were then screened by the second author for studies involving language and including brain imaging. Secondary screening was for the following criteria: 1) fMRI or PET studies (not DTI, ERP, or structural imaging) 2) whole brain activation analysis (not region-of-interest, resting state functional connectivity, dynamic causal modeling, or independent components analysis) and 3) reporting results for healthy adult participants.

### 2.2. Meta-data coding

Experimental contrasts were categorized as grammatical (rule-following) or non-grammatical (rule-violating). Meta-data coding for each paper included task definition, stimulus type, stimulus presentation modality, response modality, amount and type of training/learning, feedback type and frequency, and the experimental contrasts. Contrasts from the coded papers included directional analyses (e.g., group differences or contrast comparisons), increasing activation, decreasing activation, correlation with task performance, and conjunction analyses. Only contrasts reporting (1) increased activation during either a rule-based learning (grammatical) or non-rule-based (ungrammatical) task, or (2) experimental contrasts reporting relevant contrasts between experimental conditions (e.g., rule-based > random) in healthy adults were included. Decreasing activations were excluded. Additionally, contrasts correlating functional activation during learning with test performance were excluded. Included grammatical contrasts were those for which activation during a grammatical task (learning or test phase) was contrasted with baseline or rest; or the comparison of activation in grammatical > ungrammatical conditions.

### 2.3. Modeled activation maps

Once the relevant papers were identified and experiments/contrasts chosen, the coordinates for brain activations were extracted. Coordinate-based meta-analyses were implemented in a revised version of the Activation Likelihood Estimation (ALE) algorithm (Eickhoff et al., 2009, 2012; Turkeltaub et al., 2011), as implemented in the Neuroimaging Meta-Analysis Research Environment (NiMARE) v0.0.3, a centralized standard implementation of meta-analytic tools through Python (Salo et al., 2020). This step identified the brain regions most commonly active during statistical/implicit learning in imaging studies of normal, healthy participants. This ALE algorithm modeled foci for each contrast with a spherical Gaussian blur with full width at half maximum determined by the number of subjects included in each experiment, representing uncertainty due to within-subject and across-lab variability. Any coordinates reported using the Talairach atlas (Talairach & Tournoux, 1988) were converted to the Montreal Neurological Institute (MNI) space (Collins et al., 1994; Evans et al., 1993) using tal2icbm (Laird et al., 2010; Lancaster et al., 2007). Then, a set of modeled activation maps for each experimental contrast was generated such that each voxel’s value corresponded to its maximum probability of activation. ALE values were calculated as the voxel-wise convolution of the modeled experimental contrasts, quantifying the spatial convergence across the brain. ALE values were transformed to *p*-values using a cumulative distribution function and thresholded at *p* < 0.001. A Monte Carlo approach was implemented to correct for multiple comparisons and determined a minimal cluster size through a set of 10,000 iterations. For every iteration, randomly selected gray matter mask coordinates replaced the foci, ALE values were calculated for the randomized dataset, and those values were transformed to *p*-values, thresholded at *p <* 0.001, and the maximum size of supra-threshold clusters was recorded. The maximum cluster sizes for each iteration were used to build a null distribution that was contrasted with the original ALE map so that those larger than the cluster size in the null distribution’s 95^th^ percentile made up the family-wise error (FWE) corrected convergence maps (i.e., cluster-forming threshold at *p_voxel-level_* < 0.001 and cluster-extent threshold at *p_FWE-corrected_* < 0.05 were used to control for multiple comparisons). Surface-based and axial slice visualizations were derived with NiLearn plotting tools (Abraham et al., 2014) at https://github.com/mriedel56/surflay.git.

The first meta-analysis assessed convergence across a dataset of the pooled artificial grammar contrasts, identifying brain regions consistently active regardless of the grammaticality of the stimuli. Two additional ALE meta-analyses were then performed to identify convergence of brain activity for foci obtained from contrasts of Grammatical (rule-following; activation when the test item correctly follows the implicitly learned rule) or Ungrammatical (rule-violating; activation when the test item incorrectly follows the rule) stimuli, thus elucidating convergence of activity based on grammaticality. Conjunction analyses examined the significant convergence, or union, common between both ALE groups. Subtraction analyses of the unthresholded Grammatical ALE and unthresholded Ungrammatical ALE helped to identify brain regions that were distinctly convergent in one versus the other task type. To do this, a null distribution of the ALE difference scores was created by pseudo-randomly permuting the experimental contrasts between ALE groups, calculating voxel-level difference scores, and repeating this procedure for 10,000 iterations. Experimental contrasts were shuffled and an equal number of contrasts to those originally in the Grammatical or Ungrammatical conditions were assigned. These pseudo-ALE images were then subtracted. Voxel-level *p*-values were assigned based on a voxel’s observed difference score relative to its null distribution of pseudo-ALE difference scores (*p_FWE-corrected_* <0.05). An additional extent-threshold of 100 contiguous voxels was also applied to exclude any small regions with spurious differences (c.f., Bartley et al., 2018; Laird et al., 2005; Poudel et al., 2020).

### 2.4. Functional connectivity profiles

Once ALE clusters were identified indicating common activity across studies for the pooled Grammatical + Ungrammatical meta-analyses, their connectivity was evaluated to define subnetworks, or cliques. Connectivity profiles were computed for the ALE-derived ROIs. This step helped to identify subgroups of functionally connected brain regions. To do this, a 6-mm radius spherical seed was generated at each local maxima from each ALE cluster of the meta-analysis maps using FSL’s cluster command. Only local maxima that were at least 20-mm from another served as an ROI. These seeds were then used to identify task-independent and task-dependent connectivity between the average ROI time-course and all other brain voxels. Unthresholded MA maps for each ROI were cross correlated with each other to generate ROI x ROI correlation matrix. The same was done for unthresholded resting state functional connectivity images for each map. ROI labels were assigned using AFNI’s *whereami* command with AAL atlas labels derived from the atlas reader python package (https://github.com/miykael/atlasreader).

### 2.5. Task-independent functional connectivity using the resting-state fMRI (rsfMRI) data from the Human Connectome Project’s (HCP) Young Adult Study (Van Essen et al., 2013; S1200 Data Release)

Each ROI from the pooled ALE was used as a seed in a seed-based query of task-independent functional connectivity between the average of each ROI time course and all other brain voxels. The S1200 data release includes minimally pre-processed and denoised MRI data per the workflow detailed by Glasser and colleagues (Glasser et al., 2016). On November 12, 2019, 150 randomly selected parjcipants (mean ± SD: 28.7 ±3.9 years old) were downloaded via the HCP’s Amazon Web Services Simple Storage Solujon repository. The sample included 77 females (30.3 ±3.5 years old) and 73 males (27.1 ±3.7 years old) who provided consent through the contribujng invesjgators’ local Insjtujon Review Board or Ethics Commiree. While this age difference between biological sex was significant (*t*[149] = 5.3, *p* < 0.001), it was consistent with that noted in the full S1200 Data Release (Van Essen et al., 2013). Each HCP participant underwent T1– and T2-weighted structural imaging and four rsfMRI (15 minutes each) acquisitions on a 3T Siemens Connectome MRI scanner with a 32-channel head coil. Parameters were – structural: 0.7-mm isotropic resolution; rsfMRI: TR=720ms, TE=33.1ms, in-plane FOV=208×180mm, 72 slices, 2.0mm isotropic voxels, and multiband acceleration factor=8 (Feinberg et al., 2010). The average time course for each ROI was extracted for each participant as well as the average time course for all brain voxels. Each ROI’s separate deconvolution (FSL’s FEAT, (Jenkinson et al., 2012) included the global signal time course, which served as a regressor of no interest, and spatially smoothed with a 6mm FWHM kernel (https://github.com/NBCLab/niconn-hcp). This was done given the use of such a regressor performs better than other commonly used motion-correction strategies for HCP rsfMRI data (Burgess et al., 2016). Each participant’s ROI was then averaged across their four rsfMRI runs using a fixed-effects analysis. A rsfMRI map was then derived through a group-level, mixed-effects analysis (Woolrich et al., 2009). Gaussian Random Field theory-based maximum height nonparametric thresholding (voxel-level FWE-corrected at *p* < 0.001) was used to create spatially specific rsFC maps (c.f., Woo et al., 2014).

### 2.6. Task-dependent functional connectivity – meta-analytic connectivity modeling (MACM)

Each ROI from the pooled ALEs was also seeded in MACM analyses (https://github.com/NBCLab/niconn). These analyses reflect the coactivation patterns of spatially distinct brain regions with the seed from varied task-based neuroimaging studies and indicate the brain regions most likely to coactivate with a given ROI across tasks and behavioral domains (Laird et al., 2009). Neurosynth (Yarkoni et al., 2011) was searched for all studies reporting coordinates from each of the ALE ROIs output from NiMARE (Salimi-Khorshidi et al., 2009). A 15-mm FWHM kernel was used for all study coordinates and thresholded with a voxel-level FWE correction of *p* < 0.001 to parallel the rsFC assessments.

### 2.7. Hierarchical clustering of the pooled ALE brain regions

Hierarchical clustering of the connectivity profiles grouped the brain regions into subnetworks that were associated with grammatical/non-grammatical processes. rsFC and MACM cross-correlation matrices were calculated separately using the unthresholded connectivity maps. Three-dimensional images that represented connectivity were vectorized and concatenated creating matrix of the number of voxels (V, 2mm MNI152 template) by the number of maps (M, ROIs). Each pair-wise combination of maps was correlated (Pearson) resulting in an MxM matrix. The similarity of the task-independent and task-dependent maps was then represented in an agglomerative hierarchical cluster tree demonstrating clusters of ROIs with similar features but that were distinct across clusters. This is done using an algorithm that finds two spatially similar clusters, merges it with the two most similar ones, and continues until all are merged as measured by a standardized Euclidean distance method and Ward’s minimum variance linkage (Eickhoff et al., 2011; Timm, 2002). These clusters, or cliques, demonstrate similar task-independent connectivity in the rsfMRI data and task-dependent coactivation patterns in the MACMs; however, given the variance in imaging modality it is likely for clustering outcomes to differ. Thus, an integrated multimodal correlation matrix that combines the averaged rsFC and MACM information was created. This multimodal correlation matrix then provided the final cluster solution/groupings as in (Hill-Bowen et al., 2021).

### 2.8. Functional decoding

Functional decoding further characterized the mental operations associated with those sub-networks. To do this, the spatial correlations between the averaged and unthresholded MACM maps across ROIs was associated with each of the 1,335 terms in Neurosynth. This produced a ranked list of psychological terms that related to each cluster, thus providing a semi-quantitative interpretation of each map to the broader literature. Here, the top 10 anatomical or functional terms (removing duplicates or synonyms) that correlated with an input map were considered to represent the functional profile of the subnetwork.

## 3. Results

### 3.1. Outcomes of the literature search

The literature searches yielded over 800 papers. The title and abstract screening reduced the total to 228 papers. Of those, 88 papers were excluded for not utilizing fMRI or PET imaging, or not conducting whole brain analyses. Thirteen papers were excluded due to not including or reporting results for healthy adults. An additional 32 papers were excluded because they administered serial reaction time task (SRT) experiments that did not involve a grammar.

A final screening was completed for the remaining 150 whole-brain fMRI or PET language studies (95 identified through the literature search and 55 articles from (Tagarelli et al., 2019)) by one author (KC) to determine whether each study fit inclusion criteria. Studies including language learning which referenced rules, grammar, regularities, statistical probability, chunk strength, dependencies, and/or adjacencies were included. These studies either utilized a word learning or a grammar learning paradigm. Studies that focused on word learning were reviewed to determine whether learning occurred implicitly (no translations given), and if there was an underlying rule system. Studies that referenced explicit grammar learning were reviewed to clarify whether rules were explicitly taught (and examine the nature of feedback), and if so, they were excluded. As a result, 36 word-learning studies were excluded that either did not involve rules or included explicitly taught words. For the grammar learning studies, one study was excluded (Musso et al., 2003) and four were retained (Fletcher et al., 1999; Skosnik et al., 2002; Yang & Li, 2012; Yusa et al., 2011). Finally, studies that explicitly assessed participants’ learning of words or grammar, such as recognition and word segmentation tasks, were included since the learning was determined to be implicit. Correlational analyses between task performance and brain activity were excluded due to the small number of papers (n=5), thus excluding (Opitz & Friederici, 2003), and leaving 25 papers.

A final set of 25 fMRI articles involved data from 506 participants (238 females, 268 males) and included 25 grammatical and 14 ungrammatical experiments/contrasts (Table 1). Twenty-three studies used artificial grammars or languages and two studies used unfamiliar, non-native languages. The types of artificial grammar or languages included BROCANTO (2 papers), Brocanto2 (1 paper), finite-state grammar (10 papers using both Reber and Markovian grammars), transitional probabilities (5 papers), and the remainder included a mix of adjacent, non-adjacent, or pairwise dependencies, hierarchical rules, chunk-based rules, or phrase structure rules (7 papers). Chunk-based rules, also known as associative chunk strength, are a learning mechanism that relies on the frequency of pairs of letters that appear together to make grammaticality judgements (Meulemans & Van der Linden, 1997). Only 8 of the studies provided some sort of feedback. Fifteen of the studies presented stimuli visually, 5 auditory, and 3 auditory + visual. A subset of papers (n=12) included greater activation for ungrammatical items greater than for grammatical items and were included in a separate ALE group. See Table 1 for a summary of papers included in the meta-analysis and a summary of study characteristics. A summary of the literature search is presented as a PRISMA flowchart (Figure 2).

**Figure 2.**
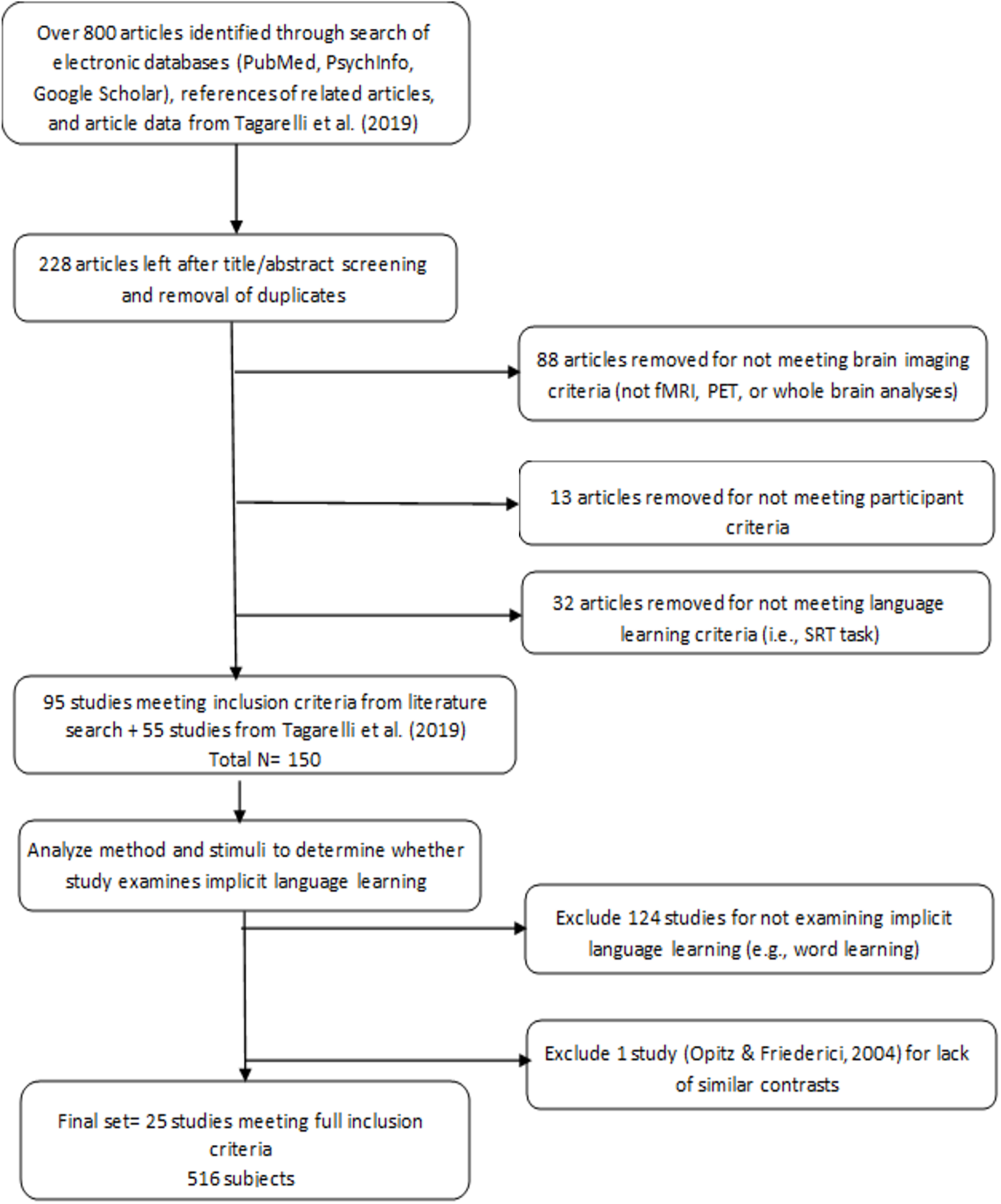
PRISMA flowchart showing the literature search and paper selection process for the meta-analysis.

**Table 1.**
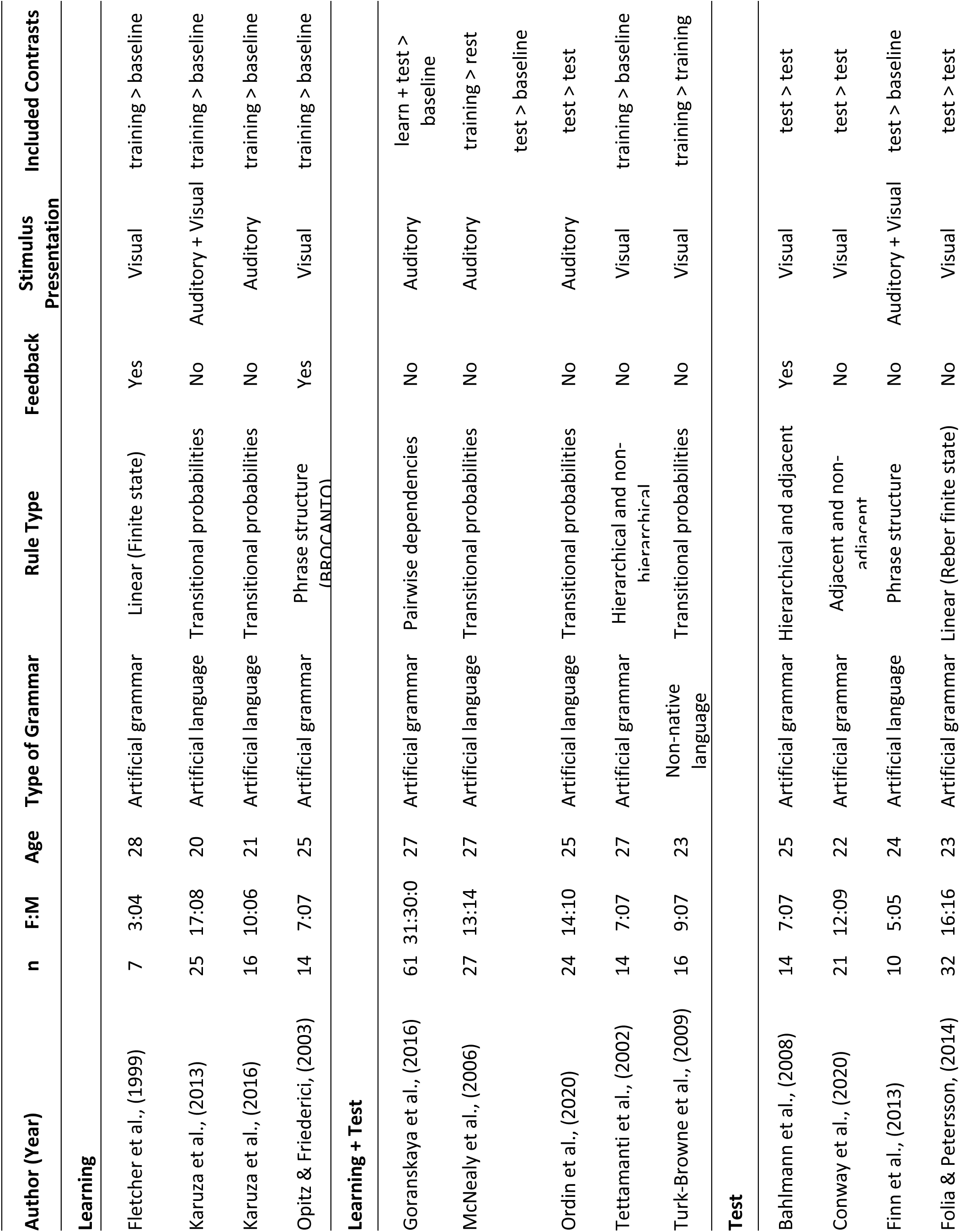
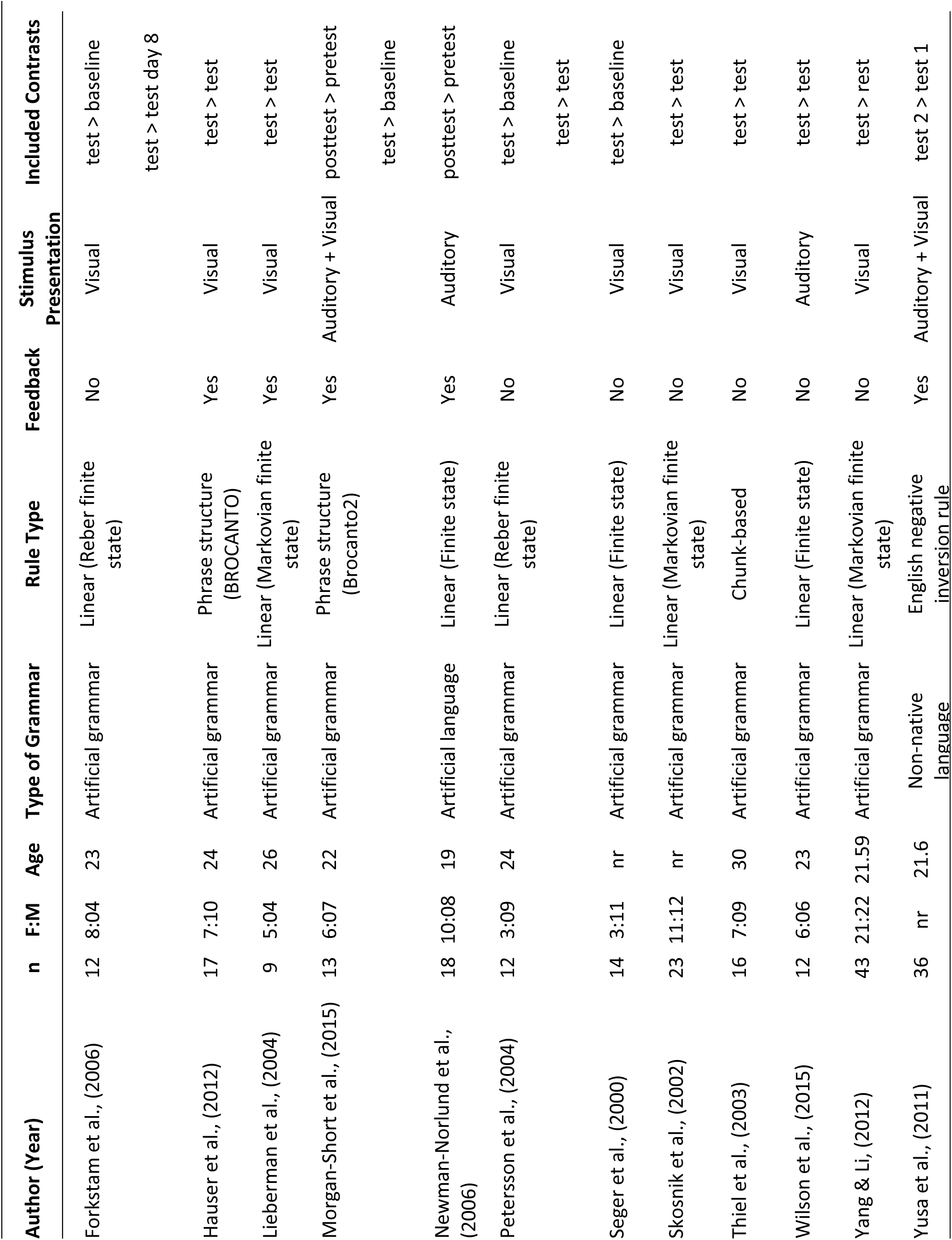
Papers included in the ALE meta-analyses are listed by condition (learning, learning + test, or test) and their study characteristics.

### 3.2. Meta-analytic outcomes

The Grammatical and Ungrammatical ALE groups were pooled to identify convergence across studies representing the recognition and endorsement/acceptance of rule-based stimuli and the rejection of random, non-rule following stimuli. The pooled ALE results identified significant activation in bilateral ROIs of the frontal, insular, and parietal cortex with six clusters of activation: left pars Opercularis extending into the left insula, right pars triangularis extending into the right precentral gyrus, right middle cingulate gyrus extending to the left supplemental motor area, right insula, right middle occipital gyrus, and right inferior parietal lobule (Figure 3A, Table 2). The statistically significant conjunction of the Grammatical and Ungrammatical ALE groups showed only three regions of convergence, organized in two main clusters including the left pars Opercularis and triangularis and the left insula (Figure 3A, Supplemental Table 1).

**Figure 3.**
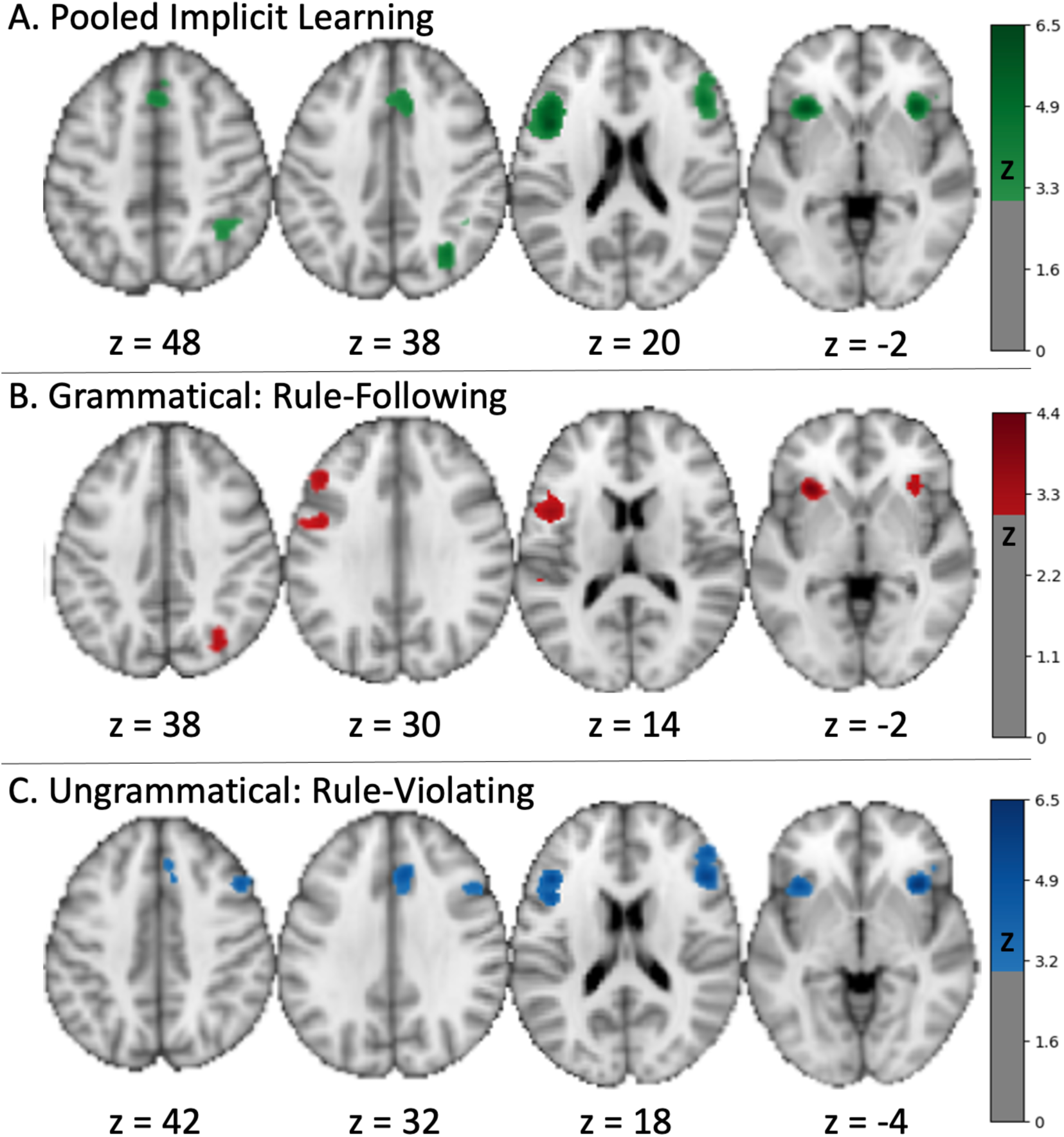
Convergent activity in the pooled, grammatical, and ungrammatical meta-analyses. The pooled (A, grammatical + ungrammatical) rule-following or rule-violating analysis identified convergent activity in bilateral inferior frontal cortex (opercular on the left, triangularis on the right), mid-cingulate to supplementary motor cortex, right mid-occipital gyrus, and right inferior parietal lobule. The convergence distinct to grammatical data (B) included a large left frontal cluster involving the opercular, triangular, and precentral cortex as well as bilateral insula and right middle occipital gyrus. Regions unique to the ungrammatical data (C) included bilateral inferior frontal, right mid-cingulate, and right insula (Supplemental Table 1).

**Table 2.** The ALE results for the pooled grammatical and ungrammatical contrasts identified 6 clusters of activity common across studies that included bilateral frontal and insular cortex along with right-sided cingulate, occipital, and parietal regions.

| *Cluster* | *x* | *y* | *z* | *Volume (mm<sup>3</sup>)* | *ALE Max* | *Label* |
| --- | --- | --- | --- | --- | --- | --- |
| 1 | -44 | 12 | 20 | 12352 | 7.36 | L Inferior Frontal Opercularis |
|  | -34 | 22 | -2 |  | 7.26 | *L Insula* |
| 2 | 48 | 26 | 20 | 5392 | 5.73 | R Inferior Frontal Triangularis |
|  | 50 | 26 | 4 |  | 4.69 | *R Inferior Frontal Triangularis* |
|  | 48 | 4 | 30 |  | 3.32 | *R Precentral Gyrus* |
| 3 | 6 | 26 | 34 | 3952 | 5.79 | R Middle Cingulate Gyrus |
|  | 2 | 22 | 50 |  | 4.99 | *L Supplemental Motor Area* |
| 4 | 36 | 22 | -4 | 2584 | 7.29 | R Insula |
| 5 | 32 | -66 | 38 | 1264 | 5.52 | R Middle Occipital Gyrus |
| 6 | 36 | -50 | 48 | 1104 | 4.02 | R Inferior Parietal Lobule |

The conjunction analysis clarified that grammatical contrasts uniquely identified five clusters comprised of seven regions across both hemispheres (Figure 3A; z corrected, FWE thresholded at *p* < 0.05; Supplemental Table 1) including the left inferior frontal gyrus extending from the precentral gyrus to the pars triangularis, left insula, right middle occipital gyrus, right insula, and left superior temporal gyrus. The ungrammatical stimuli uniquely activated four clusters comprised of eight regions (Figure 3C; z corrected, FWE thresholded at *p* < 0.05; Supplemental Table 1) in the left pars Opercularis extending to the left insula, right middle frontal gyrus extending to the right pars orbitalis, right middle cingulate gyrus extending to the left supplemental motor areas, and the right insula. However, regional activity did not significantly differ when participants were distinguishing rule-following versus rule-violating stimuli (*p_FDR_* > 0.05). Visual inspection of the maps suggests subtly more left-lateralized activation as well as left superior temporal gyrus activation when the stimuli were rule-following (grammatical).

### 3.3. Functional connectivity and hierarchical clustering outcomes

The ROIs from the pooled grammatical + ungrammatical ALE were used as seeds to assess connectivity in the rsFC and MACM data (Figure 4). Connectivity profiles indicated strong connections amongst the ROIs, as well as inclusion of subcortical structures (e.g., thalamus, basal ganglia), but the general pattern was consistent with structures of the frontoparietal network. Hierarchical clustering of the connectivity profiles, or cliques, grouped brain regions into subnetworks with processes common across spatial and functional domains. Cross-correlations of the *unthresholded* MACMs and rsFC maps for each ROI generated correlation maps that were then averaged. Hierarchical clustering was performed on each matrix and visual inspection of the metrics and dendrograms (Figure 5) indicated a of three cluster solution was optimal. Brain maps of these clusters demonstrate **considerable** overlap in frontoparietal cortex, but with considerable connectivity that includes subcortical, basal ganglia regions in one (red in Figure 5).

**Figure 4.**
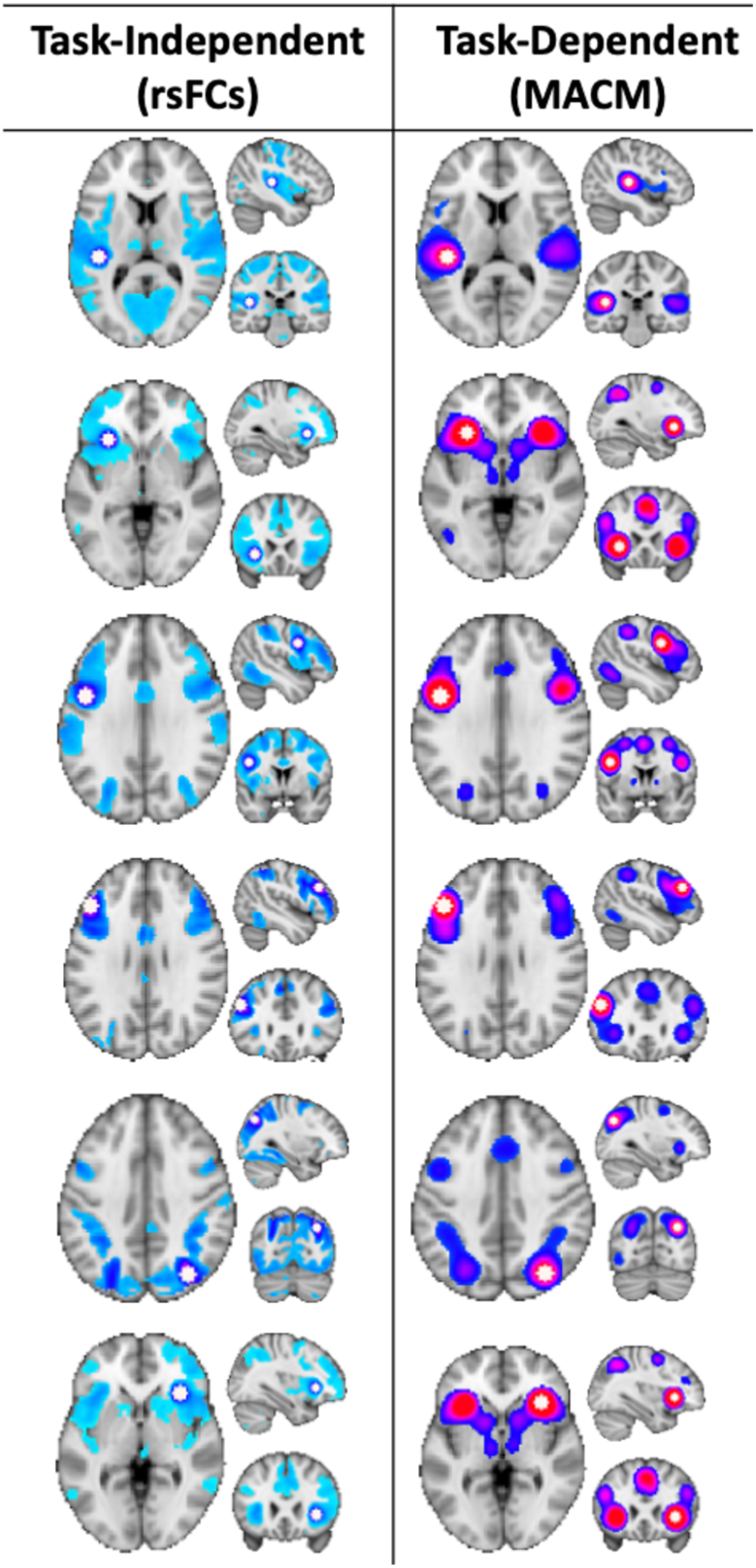
Connectivity amongst the pooled ALE-derived ROIs was characterized by using them as seeds in task-independent (resting-state functional connectivity) and task-dependent functional (meta-analytic connectivity modeling) connectivity analyses.

**Figure 5.**
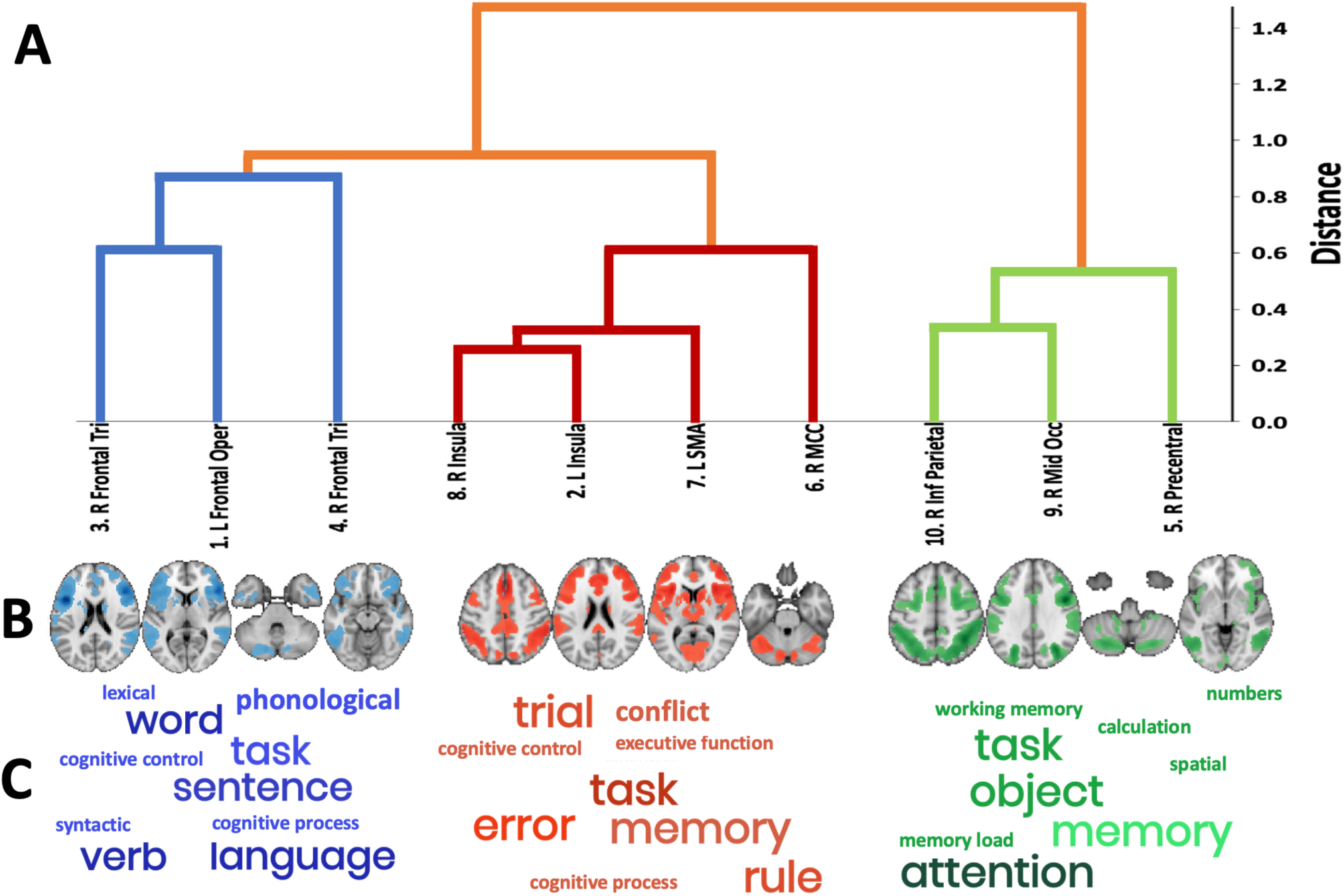
Subgroups of artificial grammar-related ROIs and their functional decoding linking them to mental operations. A) Hierarchical clustering dendrogram presenting the final sub-networks (B) based on the integrated multimodal clustering. C) Visual presentation of the clique’s and their associated terms with font size indicative of the strength of the correlation.

### 3.4. Functional decoding

Spatial correlations between the map for each of the three hierarchical clusters and the terms in the Neurosynth allowed for annotation of terms most related with activity or connectivity of each. This step distinguished functional specialization of the clusters to one clique aligned with language, one with cognitive control, and one with attention (Figure 5). Thus, the following functional interpretations of the clusters were suggested:

- Clique 1: A language (Figure 3, blue) clique was composed of the left inferior frontal Opercularis, the right inferior frontal gyrus and right inferior frontal triangularis. This clique was associated with the following Neurosynth terms: *phonological, language, word, demands, syntactic, linguistic, lexical, semantic,* and *languages*. This decoding aligns the functional specialization of this clique with processing of linguistic stimuli regardless of the correctness of grammatical rules.
- Clique 2: A salience (Figure 3, red) clique was composed of bilateral insula, right mid-cingulate cortex, and the left supplemental motor area. It is, in part, consistent with the canonical salience network (Seeley et al., 2007) or the cingulo-opercular network (Coste & Kleinschmidt, 2016). This clique associated with the Neurosynth terms *task, cognitive control, conflict, working memory, stop signal, response inhibition, demands*, and *error*. Thus, this clique is associated with the recognition of multiple features present in the stimuli and the conflict associated with determining the correctness.
- Clique 3 (Figure 3, green): A cognitive control or attentional (Figure 3, green) clique was composed of posterior regions in the right inferior parietal lobule and right mid-occipital gyrus, as well as the right precentral gyrus. This clique associated with Neurosynth terms like those of clique 2 (i.e., tasks, working memory) as well as *calculation, attentional, numbers, spatial,* and *numerical*. The decoding of this clique realized commonalities with clique 2 relative to working memory and demand, but also aligns with the executive control network (Seeley et al., 2007) or frontoparietal network (Cole et al., 2013; Cole & Schneider, 2007).

## 4. Discussion

This study presents a common set of brain regions active across many neuroimaging investigations during artificial or non-native grammar learning and their functional connectivity profiles. Using coordinate-based meta-analysis methods to identify those common regions, large task-independent and task-dependent datasets were leveraged to identify the functional connectivity profiles for these regions across the brain. From those data, subnetworks, or cliques, of regions were identified and associated with the mental operations frequently associated with them. These profiles suggest three subnetworks that are engaged for artificial and non-native grammar learning – a language network, a salience network, and a cognitive control network. The tasks included in the meta-analyses varied for modality of input (auditory, visual, or both) and thus the findings suggest that the engagement of these subnetworks for ISL is domain-general, though future analyses contrasting modality of input may further delineate the role of modality-specificity.

### 4.1 Brain Regions Commonly Active for Artificial and Non-Native Grammar Tasks

As has been reported previously in ISL tasks, the left inferior frontal gyrus was engaged for artificial and non-native grammar learning but was not uniquely involved for grammatically correct, or rule-following, stimuli. This suggests that the left IFG is necessary for both the extraction of rule governed patterns and the resolution of conflict for stimuli not following the pattern. Along with the left IFG was activity of right IFG, bilateral insula, right precentral gyrus, right mid cingulate gyrus and supplementary motor area, right middle occipital gyrus, and right inferior parietal lobule. These findings support a left-dominant, but bilateral group of regions involved largely for identification of rule-following and rule-violating grammatical stimuli, regardless of the modality of the input. Notably missing from the meta-analysis results were the basal ganglia, given their proposed primary role in ISL. This is somewhat consistent with the meta-analytic findings of Tagarelli and colleagues (2019), but they also investigated common activity in lexical learning and in non-declarative grammar learning where they found left caudate and putamen activity. Plante and colleagues (2015) also found caudate activity, but only in the early phases of learning. Both Tagarelli et al. (2019) and Plante et al. (2015) note that caudate activity was specific to grammar learning and particularly in the early extraction of patterns or rules in the stimuli. As such, it may be that the collapsing across learning phases specific to rule-learning precluded the finding in the present study.

### 4.2 Functional Connectivity Profiles

Utilizing the regions that were commonly active in the grammar learning tasks as seeds to query connectivity in resting-state and task-dependent data sets, profiles of connectivity elucidated three separate subnetworks. These three networks were further characterized through functional decoding to clarify that one was likely specialized in language processing, one in salience of features in stimuli, and one in cognitive control (Figure 3).

These profiles extend from the specific regions active during the task to include a more diffuse representation of each region’s participation in brain networks, thus providing evidence to build hypotheses for how these networks may contribute to task performance. For example, the left inferior frontal gyrus (pars Opercularis) that is proposed to be central to grammar learning was found to be commonly active across the artificial grammar learning studies, and it is part of a network of brain regions (clique 1, blue in Figure 3), including the right inferior frontal gyrus (pars triangularis), and the left insula. This network of regions associated with terms in NeuroSynth that clarified its role in language with specific terms like phonological, syntactic, semantic, sentence, as well as to a lesser extent in general cognitive processing with specific terms including demands, domain-general, working memory, and cognitive control.

While this network includes the left IFG, there is an equal or greater extent of right IFG inclusion in the network, suggesting that grammar learning utilizing auditory and visual stimuli may not be uniquely left dominant or language centric.

While the basal ganglia (specifically the bilateral caudate nuclei) were not commonly active across the studies included in the meta-analyses, they were identified as part of clique 3, a central executive, frontoparietal network. Prior evidence suggests that this cognitive control network flexibly shifts as the cognitive goals or demands of a given task change (Cole et al., 2013). And, as noted above, the caudate nucleus has been found to be active during the early phases of grammar learning (Plante et al., 2015; Tagarelli et al., 2019), but not once the grammatical rule has been learned. Thus, we propose that clique 3 is involved in grammar learning as a scaffold for the changing demands in learning across grammar rule identification, maintenance, and rule application in healthy adults.

In concert with the network involving the left IFG (clique 1), and the network involving the basal ganglia (clique 3), another network engaged for cognitive control included frontoparietal regions that associated with behaviors like working memory and attention (clique 2). This network clustered with the insula and mid-cingulate cortex regions derived from the ALEs and we propose its role in ISL is for realizing the salience of features in the stimuli and their relevance in rule-learning. This network would be necessary to track regularities across stimuli and imperative for rule identification and application of grammar, as has been predicted by others reporting frontoparietal activity during language learning tasks. The frontoparietal brain network is domain-general and largely overlaps with other cognitive control networks (e.g., multi-demand), the integrity of which is known to be a positive prognostic sign for language recovery in individuals with aphasia (Fedorenko et al., 2013). Future study will determine whether connectivity of the additional subcortical structures, and other temporal structures, are also relevant to successful language learning.

### 4.3. Limitations

The approach taken and interpretations made in this study are based on meta-analytic methods from large-scale data mining. The ALE conducted to identify regions of interest was limited to functional activation studies utilizing similar artificial grammar learning tasks. This approach is susceptible to publication bias. That is, it is limited by the design and reporting of results in the primary published studies. As well, the coordinate-based ALE algorithm does not take into account the cluster size of the original study, which is likely less precise than results of an image-based meta-analysis (Salimi-Khorshidi et al., 2009).

The 25 artificial language/grammar studies included in the meta-analysis were common in their inclusion of an implicit-statistical learning paradigm and contrasts of rule-following or rule-violating stimuli. However, the studies differed in terms of the grammatical rule types, the inclusion of feedback during learning trials, and whether the stimuli were auditory (n=6), visual (n=15), or both (n=4) (see Table 1). The common clustered activity across studies is the primary outcome of the ALE, but we recognize that there may be bias toward the study types with more representation – finite state artificial grammars learned without feedback and presented in visual stimuli. Future study may focus on artificial grammars with hierarchical structures that more closely resemble natural language, and which may expand the set of ROIs and continue to clarify the specificity or interactions amongst the subnetworks identified here when grammars are more complex.

## 5. Conclusions

By utilizing advanced neuroimaging meta-analytic techniques, this study elucidates the brain regions actively engaged during artificial grammar learning tasks. These included the oft-hypothesized left IFG as well as other frontal, insular, cingulate, and posterior parieto-occipital cortices, consistent with findings of other meta-analytic work. By seeding these regions in large task-dependent and task-independent datasets and applying hierarchical clustering, functionally connected subnetworks were identified. These subnetworks were then linked to mental operations commonly attributed to their collective activity. In this way, three subnetworks, or cliques, were found and suggest bilateral connectivity profiles engaged for language, salience, and cognitive control. The language subnetwork (clique 1 in Figure 5) was not left-dominant, though the ALE results contrasting rule-following and rule-violating stimuli may indicate more left-sided activity for grammatical correctness. The other two subnetworks had bilateral representation and their association with mental operations suggest they are engaged for domain-general processing.

These findings provide a brain network model of ISL for artificial and non-native grammars.

This model suggests recruitment of a left hemisphere dominant language network as well as bilateral salience and cognitive control networks for learning rule-governed stimuli. The model thus presents three differentiated brain networks integral to learning, though the specific role that each play in successful learning requires further investigation. As noted in the Introduction, it is likely that the language network is damaged in individuals with aphasia, but the integrity of the cognitive control and salience networks may be relatively spared. And, if they are spared then there is evidence of better prognosis (Fedorenko et al., 2013; Geranmayeh et al., 2014). If it is the case that one of these networks is impaired in a clinical population, like aphasia, then understanding whether designing intervention approaches that optimize the function of one of the other networks may be key to improving learning, and thereby the therapeutic outcomes.

## 6. Data and code Availability

The data and code supporting the findings of this study are available on Github (https://github.com/NBCLab/meta-analysis_implicit-learning) and on NeuroVault (https://neurovault.org/collections/9597/). Tools used to perform the rsFC analyses can be found in these GitHub repos: https://github.com/NBCLab/niconn; https://github.com/NBCLab/niconn-hcp. Code for plotting of the results is at Github (https://github.com/mriedel56/surflay).

## 7. Author Contributions

Conceptualization, A.E.R. and K.C., Methodology: M.C.R., A.R.L, and A.E.R., Formal Analysis: M.C.R., K.C., and A.R.L., Writing, A.E.R, K.C., J.C.T., K.L., and A.R.L., Visualization, M.C.R. and A.E.R., Supervision, A.E.R. and A.R.L.

