## Supplemental Table 1 for "Elucidating an implicit-statistical learning brain network: Coordinate-based meta-analyses and functional connectivity profiles of artificial grammar learning in healthy adults"

| ALE GROUP | CLUSTER | X | Y | Z | Volume ($mm^3$) | ALE Max | Label |
|---|---|---|---|---|---|---|---|
| ***GRAMMATICAL*** | | | | | | | |
| | 1 | -46 | 8 | 14 | 6096 | 5.08 | L Inferior Frontal Opercularis |
| | | *-46* | *28* | *28* | | *4.84* | *L Inferior Frontal Triangularis* |
| | | *-46* | *2* | *30* | | *4.15* | *L Precentral Gyrus* |
| | 2 | -30 | 22 | -2 | 1536 | 6.45 | L Insula |
| | 3 | 32 | -72 | 38 | 976 | 4.47 | R Middle Occipital Gyrus |
| | 4 | 34 | 24 | 0 | 888 | 4.63 | R Insula |
| | 5 | -42 | -28 | 10 | 832 | 4.55 | L Superior Temporal Gyrus |
| ***UNGRAMMATICAL*** | | | | | | | |
| | 1 | -44 | 12 | 22 | 7504 | 6.56 | L Inferior Frontal Opercularis |
| | | *-38* | *20* | *0* | | *5.78* | *L Insula* |
| | 2 | 48 | 26 | 18 | 6560 | 6.21 | R Inferior Frontal Triangularis |
| | | *48* | *18* | *42* | | *4.88* | *R Middle Frontal Gyrus* |
| | | *46* | *32* | *-4* | | *3.33* | *R Inferior Frontal Orbitalis* |
| | 3 | 6 | 26 | 32 | 3224 | 5.31 | R Middle Cingulate Gyrus |
| | | *0* | *24* | *52* | *3224* | *4.40* | *L Supplemental Motor Area* |
| | 4 | 36 | 22 | -4 | 1856 | 7.00 | R Insula |
| ***CONJUNCTION*** | | | | | | | |
| | 1 | -44 | 10 | 18 | 1856 | 4.74 | L Inferior Frontal Opercularis |
| | | *-44* | *26* | *22* | | *3.74* | *L Inferior Frontal Triangularis* |
| | 2 | -34 | 20 | -2 | 520 | 5.02 | L Insula |

**Supplemental Table 1.** Results for the grammatical and ungrammatical ALE contrasts and their conjunction.
